# Environmental DNA facilitates accurate, indigenous-led, inexpensive, and multi-year population estimates of millions of Eulachon fish

**DOI:** 10.1101/620500

**Authors:** Meredith Pochardt, Jennifer M. Allen, Ted Hart, Sophie D. L. Miller, Douglas W. Yu, Taal Levi

## Abstract

Although environmental DNA shed from an organism is now widely used for species detection in a wide variety of contexts, mobilizing environmental DNA for management requires estimation of population size and trends rather than simply assessing presence or absence. However, the efficacy of environmental-DNA-based indices of abundance for long-term population monitoring have not yet been assessed. Here we report on the relationship between six years of mark-recapture population estimates for eulachon (*Thaleichthys pacificus*) and ‘eDNA rates,’ which are calculated from the product of stream flow and DNA concentration. Eulachon are a culturally and biologically important anadromous fish that have significantly declined in the southern part of their range but were historically rendered into oil and traded. Both the peak eDNA rate and the area under the curve of the daily eDNA rate were highly predictive of the mark-recapture population estimate, explaining 84.96% and 92.53% of the deviance respectively. Even in the absence of flow correction, the peak of the daily eDNA concentration explained an astonishing 89.53% while the area under the curve explained 90.74% of the deviance. These results support the use of eDNA to monitor eulachon population trends and represent a >80% cost savings over mark-recapture, which could be further increased with automated water sampling, reduced replication, and focused temporal sampling. Due to its logistical ease and affordability, eDNA sampling can facilitate monitoring a larger number of rivers and in remote locations where mark-recapture is infeasible.

## Introduction

While the environmental DNA shed from an organism is now widely used for species detection in a wide variety of contexts (Barnes & Turner, 2016; Rees, Maddison, Middleditch, Patmore, & Gough, 2014; Jerde, Mahon, Chadderton, & Lodge, 2011; Laramie, Pilliod, & Goldberg, 2015; Mächler, Deiner, Steinmann, & Altermatt, 2014; Rees et al., 2014; Takahara, Minamoto, Yamanaka, Doi, & Kawabata, 2012), mobilizing environmental DNA for management requires estimation of population size and trends rather than simply assessing presence or absence. Recent research suggests that eDNA quantified with real-time quantitative polymerase chain reaction (PCR) or digital-droplet PCR can provide a proxy for actual abundance in controlled experiments (Rees, Maddison, Middleditch, Patmore, & Gough, 2014), in ponds (Lacoursière-Roussel, Côté, Leclerc, & Bernatchez, 2016; Takahara et al., 2012) in streams (Doi et al., 2015; Levi et al., 2019; Lodge et al., 2012; Tillotson et al., 2018; Wilcox et al., 2016) and in marine bays (Plough et al., 2018) However, the efficacy of environmental DNA based indices of abundance in natural settings have produced mixed results (Yates, Fraser, & Derry, 2019) and have not yet been assessed in a management context for long-term population monitoring.

Anadromous fish enter freshwater systems to spawn, often in large number, representing the possibility to quantify the size of the spawning population with environmental DNA to inform management and population trends. While recent research has suggested that daily eDNA counts correlate well with daily entry by adult salmon or daily outmigration of salmon smolts (Levi et al., 2019), a more important question is whether total run sizes can be accurately predicted for long interannual population monitoring programs.

Owing to their short run time and large spawning aggregations, Eulachon (*Thaleichthys pacificus*), a lipid-rich, anadromous smelt of the family Osmeridae (Mecklenburg et al. 2002), make an ideal case study to test eDNA for long-term population monitoring of anadromous fish. Adult eulachon have an average size of 18 to 22 cm (Spangler, 2002). The historic range of eulachon stretched from southern California to the Bering Sea in southwest Alaska (Hart, 1973). The majority of eulachon populations have been declining since the 1990s (Hay & Mccarter, 2000). In 2010, the National Marine Fisheries Service (NMFS) listed the southern distinct population segment in Washington, Oregon, and California as Threatened under the Endangered Species Act (NOAA, 2010). Because there is no commercial eulachon fishery in northern Southeast Alaska, there is no harvest regulation or management, agency oversight, or monitoring of population trends. While some eulachon population declines have been well documented (Hay & Mccarter, 2000), the status of most eulachon populations is either unknown or anecdotal.

In Southeast Alaska, they are the first anadromous fish to return after the long winter, and as a result, are a key resource for indigenous people and for wildlife. For the Northwest Coast native people, eulachon are a culturally significant staple food source that is consumed fresh, dried, or smoked, and are frequently rendered into oil (Betts, 1994). Historically, eulachon oil was the most important trade item on a network of ‘grease trails’ between coastal and interior peoples, and it is still used and traded (Betts, 1994; Moody & Pitcher, 2010). Eulachon spawn just prior to the breeding season of many consumers, including marine mammals, thus providing a high-energy prey resource at an energetically demanding time (Sigler, Womble, & Vollenweider, 2004). The eulachon spawning aggregation draws enormous congregations of seabirds, bald eagles (*Haliaeetus leucocephalus*), Steller sea lions (*Eumetopias jubatus*), harbor seals (*Phoca vitulina*), and humpback whales (*Megaptera novaeangliae*) among many other smaller predators and scavengers. A lack of eulachon population information and the cultural and subsistence value of the species led to the development of an indigenous-led eulachon monitoring program in northern Southeast Alaska. In 2010 the Chilkoot Indian Association and the Takshanuk Watershed Council initiated a modified Lincoln-Petersen (Chapman, 1951, Lincoln, 1930; Petersen, 1896) mark-recapture population estimate on the Chilkoot River near Haines, Alaska at the northern end of southeast Alaska (Fig. 1). This program was successful in gathering baseline eulachon population data where none existed previously; however, monitoring is challenging and expensive (∼$20,000 annually), limiting the feasibility of conducting long-term monitoring and limiting the possible geographic scope of monitoring. In an effort to develop a more cost-effective monitoring method, in 2014 we began pairing the mark-recapture program with daily water sampling to evaluate the efficacy of environmental DNA (eDNA) to produce an index of eulachon abundance.

**Figure 1.**
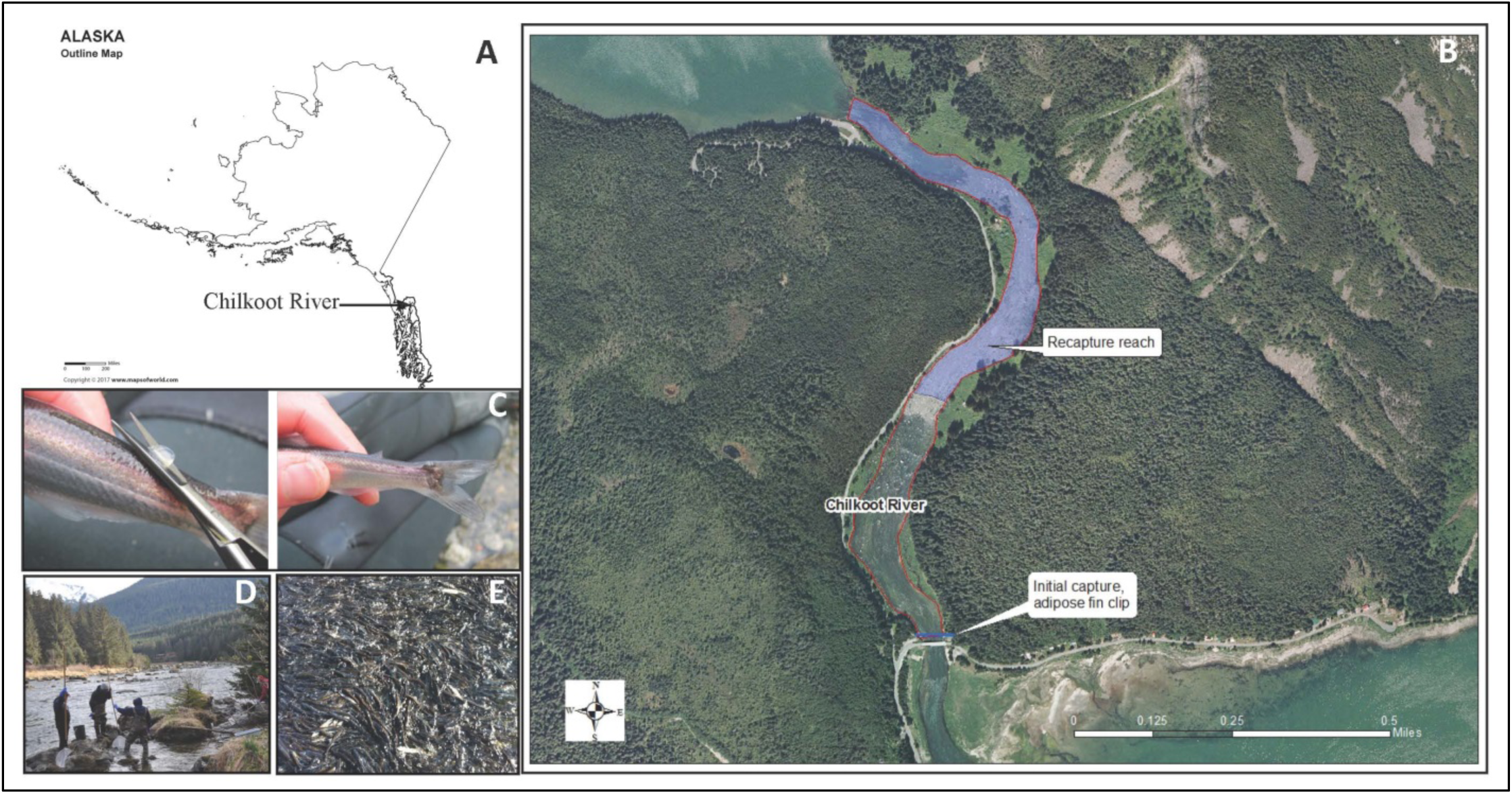
The Chilkoot River (A) is located in northern Southeast Alaska. The study area is located in the approximately 1.75 km section of river between the outlet of Chilkoot Lake and where the river meets the estuary (B). Three 1L water samples were collected at low tide just below the initial capture location, where (C) the adipose fin of eulachon captured in the trap was clipped into a “shark fin” for easy identification. The beginning of the recapture reach is located approximately 0.75 km upstream of the initial capture location, although eulachon are not exclusively recaptured within this reach. (D) Crews dip net or cast net to capture eulachon. (E) Depicts the Chilkoot River during a large eulachon run.

Here we compare six years (2014-2019) of mark-recapture eulachon abundance estimates with eulachon eDNA quantification to test whether long-term, affordable, indigenous-led monitoring of eulachon populations could be effectively achieved with environmental DNA. This method could facilitate intertribal cooperation for affordable monitoring of a culturally important subsistence and economic resource on a regional scale. Such regional monitoring is particularly important for eulachon, which exhibit low site-fidelity and thus regional broad-scale population structure (Flannery, Spangler, Norcross, Lewis, & Wenburg, 2013) such that a true population decline can only be verified by monitoring multiple river systems.

## Methods

### Study System

The Chilkoot River near Haines, Alaska has long been a culturally and ecologically important river. The lower Chilkoot River flows 1.5 km from Chilkoot Lake to the ocean at the terminus of a large fjord. The Chilkoot Tlingit village and fishcamp was historically located along the banks of the Chilkoot River, which is still utilized for eulachon fishing and processing today (Betts, 1994; Olds, 2016). Eulachon typically spawn in the lower reaches of the Chilkoot River (Hay & Mccarter, 2000) where mostly indigenous harvesters capture large quantities for smoking, frying, and rendering into oil in pits.

The Chilkoot Indian Association initiated a eulachon mark-recapture study to develop the first population baseline for the Chilkoot River, which is now the longest eulachon population dataset in Southeast Alaska (Alaska Department of Fish and Game Aquatic Resources Permit: SF-2014-027, SF2015-066, SF2016-113, SF2017-062, SF2018-072). This effort was initiated because anecdotal observation suggested that the run size and timing on the Chilkoot River differed from traditional knowledge, and because the decline of the southern distinct population segment of eulachon was substantial enough to warrant threatened status under the Endangered Species Act (NOAA, 2010). The Endangered Species Act listing of eulachon led to concern by Chilkoot Indian Association tribal members that a decline in northern Southeast Alaska, where a strong subsistence fishery remained, would go undocumented, and thus un-remediated, without quantification of the current run size (Olds, 2016).

### Mark-Recapture

At the mouth of the Chilkoot River, eulachon were captured using a modified fyke net trap and dip nets. The initial captured eulachon (M group) were transferred in small groups to plastic dishpans where they could be easily handled to clip off the adipose fin using retina scissors and returned to the river. To avoid excessive increases in temperature and to reduce the possibility of disease transmission, the water in the dishpans was changed between each group and the dishpans were rinsed with river water. To allow time for the marked fish to mix with the unmarked fish, the recapture group was captured approximately 0.75 km upstream of the trap location (C and R group) (Fig 1). Eulachon in the second capture group were collected by field crews wading through the river with dip nets making sure to sample all portions of the river and with the help of subsistence harvesters. The captured fish were examined for a clipped adipose fin before releasing. To avoid repetitive sampling of the same fish, field crews started at a downstream point and worked their way upstream. Eulachon are thought to be semelparous (spawning only once), which negates recapturing fish marked in a previous year (Clarke, Lewis, Telmer, & Shrimpton, 2007). A modified Lincoln-Peterson estimator equation (Chapman, 1951) was used 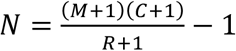 where N = total population size, M = marked initially, C = total in second sample, and R = marked recaptures. The standard error was calculated using the equation 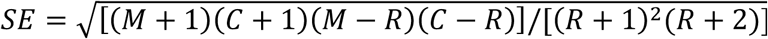. The 95% confidence interval was calculated as *CI* = *N* ±(1.96)(*SE*). Mark-recapture data were collected from 2010 through 2019, excluding 2013 where a lapse in funding prohibited collection.

### Environmental DNA

We collected daily water samples for eulachon eDNA quantification just below the mark-recapture trap location near the mouth of the Chilkoot River (Fig. 1) from 2014 through 2019. The samples were taken as close to low tide as was feasible to avoid either DNA intrusion from the estuary and/or dilution with an influx of tidal flow. Three replicate 1 L water samples were collected from the same location each sampling day in sterile Whirl Pak bags starting in early to mid-April and continuing for at least one week beyond the end of the mark-recapture study duration (Table 2). The exception to this was 5 days in 2019 for which field crews mistakenly filtered only 750 ml. We multiplied DNA concentrations from these days by 1.33 to account for the reduced volume. We sampled for 8, 11, 19, 13, 17, and 25 days during each run from 2014 through 2019 respectively.

**Table 1.** Annual eulachon population estimates for the Chilkoot River using a modified Lincoln-Peterson equation (excluding 2013). eDNA monitoring began in 2014.

| Measurement | 2010 | 2011 | 2012 | 2014 | 2015 | 2016 | 2017 | 2018 | 2019 |
| --- | --- | --- | --- | --- | --- | --- | --- | --- | --- |
| **M = Marked Initially-adipose clipped** | **8,017** | **49,814** | **27,525** | **24,084** | **306** | **9,384** | **33,681** | **30,542** | **70,127** |
| **C = Total in second sample captured above weir** | **20,210** | **143,444** | **48,376** | **19,886** | **3,122** | **8,865** | **47,654** | **18,636** | **80,859** |
| **R = Marked recaptures above weir with clip** | **72** | **568** | **186** | **140** | **2** | **45** | **126** | **64** | **210** |
| **N1 = Population Estimate** | **2.2 Million** | **12.6 Million** | **7.1 Million** | **3.4 Million** | **319,586** | **1.8 Million** | **12.6 Million** | **8.7 Million** | **26.7 Million** |
| **SE<sup>2</sup> = Standard Error** | **256,415** | **521,961** | **516,583** | **283,226** | **158,934** | **262,518** | **1,113,520** | **1,074,932** | **1,840,573** |
| **CI<sup>3</sup> = 95% Confidence Interval** | **1.7 to 2.7 Million** | **11.5 to 13.6 Million** | **6.1 to 8.1 Million** | **2.9 to 3.9 Million** | **8,074 to 631,098** | **2.3 to 1.3 Million** | **10.5 to 14.8 Million** | **6.6 to 10.9 Million** | **23.2 to 30.4 Million** |

**Table 2:** Annual field effort for the mark-recapture study and number of eDNA sample days, water samples, and ddPCR replicates.

| Year | Start Date | End Date | Number of Mark-Recapture field days | Number of eDNA sample days | Number of water samples | Number of ddPCR replicates |
| --- | --- | --- | --- | --- | --- | --- |
| 2010 | 4/23/2010 | 4/27/2010 | 5 | NA | NA | NA |
| 2011 | 4/27/2011 | 5/8/2011 | 12 | NA | NA | NA |
| 2012 | 5/2/2012 | 5/7/2012 | 6 | NA | NA | NA |
| 2014 | 5/5/2014 | 5/9/2014 | 5 | 8 | 24 | 48 |
| 2015 | 4/26/2015 | 4/29/2015 | 4 | 11 | 33 | 66 |
| 2016 | 4/20/2016 | 4/24/2016 | 4 | 19 | 57 | 114 |
| 2017 | 4/28/2017 | 5/5/2017 | 8 | 13 | 39 | 78 |
| 2018 | 5/6/2018 | 5/11/2018 | 6 | 17 | 51 | 102 |
| 2019 | 4/25/2019 | 5/7/2019 | 13 | 25 | 75 | 150 |

Each sample was transported from the field to the Takshanuk Watershed Council office immediately after collection and was filtered through a Nalgene 47mm 0.45 micron cellulose nitrate filter using either a peristaltic pump (Proactive Alexis peristaltic pump) or vacuum pump (Gast model DOA-P704-AA) with a three-sample manifold. Filters were stored immediately in 100% ethanol within 2 mL cryovials and refrigerated until shipped to Oregon State University for extraction. Filters were removed from ethanol and air-dried overnight in sterile, disposable weight boats in a hepafiltered and UV-irradiated cabinet within a PCR-free laboratory to avoid contamination. DNA was then extracted using the Qiagen DNeasy Blood and Tissue kit modified to include a >48 hour soak in ATL buffer, which was found to produce higher and more consistent yields. DNA was eluted in a total volume of 100 μl.

### DNA Quantification

We developed a species-specific quantitative PCR assay for eulachon targeting a 187-bp region of the *Cytochrome oxidase I* (COI) region of the mitochondrial genome based on observed sequence divergence among Osmeridae fish species in the Pacific Northwest region of North America including longfin smelt (*Spirinchus thaleichthys*), capelin (*Mallotus villosus*), and rainbow smelt (*Osmerus mordax*). Specifically, we ensured at least 2 bp mismatch on the forward primer and at least 3 bp mismatches on the probe to the other Osmeridae fishes. The reverse primer contained a 2 bp mismatch to longfin smelt, a 3 bp mismatch to capelin, and a 1bp mismatch to rainbow smelt. We tested our primers *in vitro* against longfin smelt tissue to ensure no nonspecific binding, and *in natura* on water samples from a diversity of rivers in southeast Alaska (Chilkoot, Chilkat, Taiya, Ferebee, Katzehin, Auke, Berners, Lace, Antler, Mendenhall) and Oregon (Columbia, Cowlitz) outside of the eulachon run to ensure no nonspecific binding to non-Osmeridae fishes.

The probe was labeled with a 5’ FAM fluorescent marker and a minor-groove-binding non-fluorescent quencher on the 3’ end. Primer3 software (Untergasser et al. 2012) was used to select the following primers: Euc_COI_R (5’-CTCCCTCCTTCCTTCTCCTT-3’), Euc_COI_R (5’-GGTCTGGTACTGGGAAATGG-3’) and the internal probe Euc_COI_I (5’-6FAM*AGCGGGAGCCGGGACTGGCT*MGBNFQ).

A Bio-Rad QX200 AutoDG Droplet Digital PCR system (Hercules, CA. USA) at the Oregon State University Center for Genome Research and Biocomputing was used to quantify DNA concentrations in duplicate. A 22 μl reaction was carried out containing (final concentrations) 1 x ddPCR Supermix for probes (no dUTP), 900 nM of both forward and reverse primers, 250 nM internal probe and 4 μl of DNA extract. Droplets were generated using the QX200 AutoDG system. Cycling consisted of 95 °C for 10 mins, followed by 45 cycles of 94 °C for 30 secs, and 60 °C for 1 min, ending with 96 °C for 10 mins, allowing for a ramp rate of 2 C/sec between steps. PCR setup occurred in a hepafiltered and UV-irradiated cabinet within a PCR-free laboratory to avoid contamination. After the reaction, the droplets were read on a Droplet Reader and analyzed with QuantaSoft Analysis Pro software (version 1.0.596). We included extraction blanks every 35 samples, and every ddPCR plate included two no-template controls (DI water), and positive controls (eulachon tissue extracts). We did not observe false positives of eulachon in negative controls nor false negatives of eulachon tissue or water samples when eulachon were observed in the river.

The concentration of eDNA is a function of both the amount of eDNA shed into the river and dilution of eDNA due to increased stream flow. To calculate the flow-corrected eDNA rate, we multiplied each day’s ddPCR DNA concentration 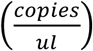 against the day’s stream flow 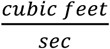. We refer to this as an eDNA rate because once the volume units cancel the result is proportional to DNA copies/second (Levi et al. 2019). Stream flow measurements were taken each day that an eDNA sample was collected immediately following the collection of the eDNA sample. To measure streamflow, we used a rating curve developed by the Alaska Department of Fish and Game for the Chilkoot River. To validate this rating curve, a stream flow measurement was taken at the beginning of each field season on the Chilkoot following the USGS velocity-area method using a type AA current meter (Turnipseed & Sauer, 2010). Following the initial calibration of the rating curve, the daily river height was measured in feet off of an established benchmark using surveying equipment, which was then transformed into a river discharge based on the rating curve (Sowa, 2015).

### Analysis

We evaluated the flow-corrected eDNA rate as an index of eulachon abundance based on two metrics. First, we use the maximum eDNA rate (i.e. size of peak). Second, we used area under the curve of the eDNA rate throughout the duration of the run. In each case, the daily eDNA concentration was the average of 6 replicates (2 ddPCR within 3 replicate water samples). DNA concentration Particularly in cases of multimodal runs, the area under the curve was expected to provide a more accurate representation of the overall biomass. We computed the area under the curve with the AUC function in the *DescTools* package version 0.99.27 (Signorell, 2019) in RStudio version 1.1.383 (RStudio Team, 2015). We additionally assessed the need for flow correction by evaluating the relationship between uncorrected eDNA concentrations using the same two metrics and mark-recapture population estimates. We used quasipoisson regression to model the mark-recapture population estimates as a function of the natural log of the two measures of eDNA rate and the uncorrected eDNA concentration.

## Results

### Mark-Recapture

The mark-recapture population estimate was initiated in 2010 and continued annually through 2019, excluding 2013 due to funding constraints. Eulachon exhibited substantial population fluctuations with a potential 5-6-year cyclic pattern for large returns (Fig. 2). The average eulachon population estimate for the mark-recapture method from 2010-2019 (excluding 2013) was 8.4 million, with a maximum of 26.7 million in 2019 (±1,840,573), and a minimum of 319,568 in 2015 (±158,934) (Table 1). Eulachon arrival in the Chilkoot River was documented as early as April 20^th^ (2016) and as late as May 6^th^ (2017), with run durations lasting between 4 (2015) and 13 days (2019).

**Figure 2.**
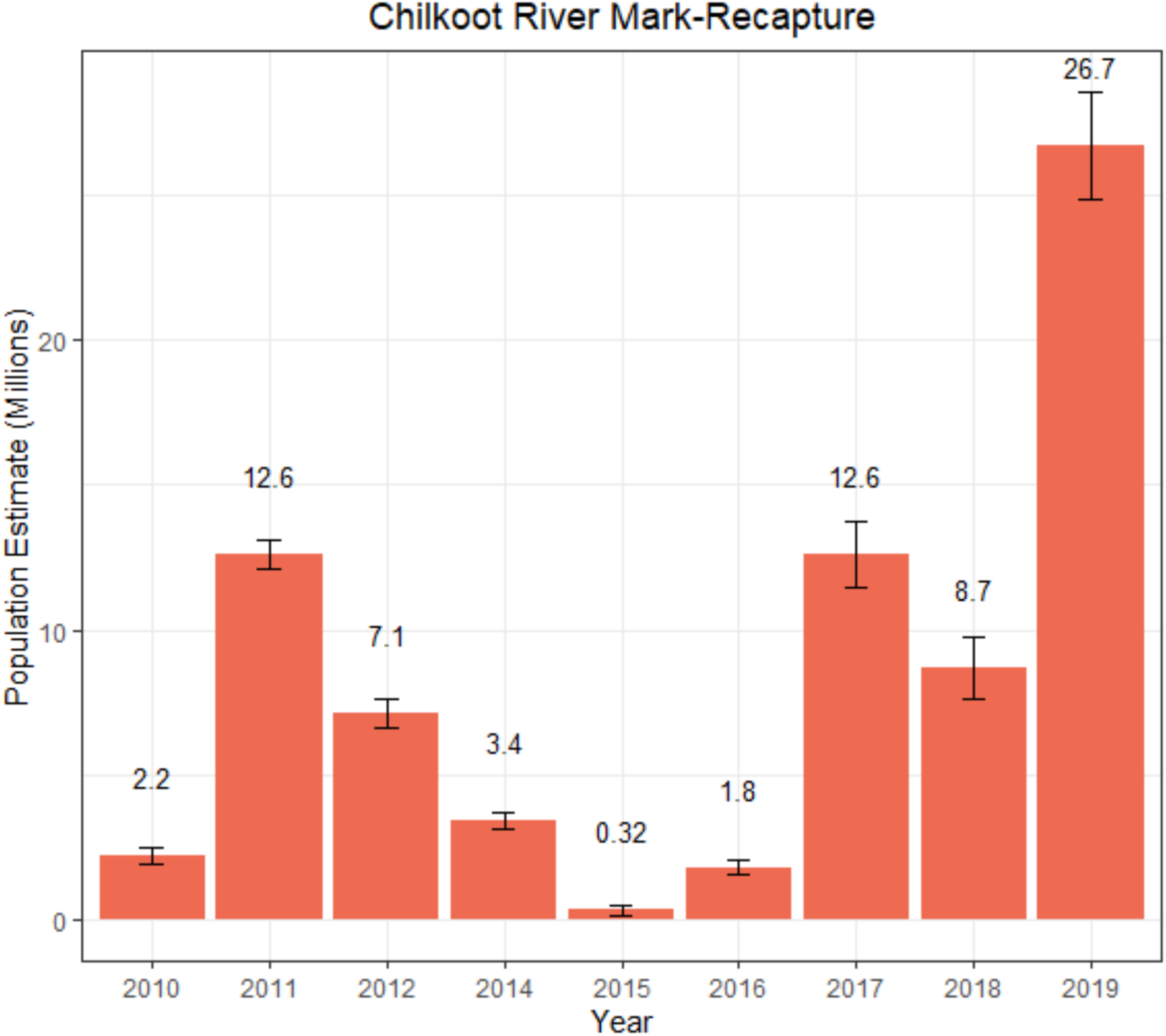
Results of mark-recapture population estimate for eulachon on the Chilkoot River using a modified Lincoln-Petersen method. Error bars represent one standard error.

Two notable anomalies occurred during the mark-recapture study period. In 2016 the run consisted of multiple pulses with what appeared to be a definitive end of the run that was followed by a final pulse of fish five days later. This final pulse of fish was not recorded as part of the mark-recapture estimate but was captured in the eDNA data. Additionally, the 2015 run was a failure, with the smallest number of fish during our study period, but anecdotal reports suggested that other rivers in Northern Southeast Alaska received unusually large runs.

### Environmental DNA

During eulachon runs, all ddPCR replicates of all technical replicates amplified with the exception of the period that appeared in the field to be prior to any obvious eulachon entry in which we either observed no amplification or very low copy number amplification of one replicate but not both. eDNA concentrations varied substantially from near zero to a high of 328000 copies / μL during the peak of the run in 2017. The product of streamflow and eDNA concentration, which we refer to as ‘flow-corrected eDNA rate’ (Fig. 3, see also Levi et al. 2019), was highly predictive of the eulachon population estimate generated through the mark-recapture method. The natural log of the eDNA peak was significantly related to, and explained 84.96% of the deviance in, the mark-recapture population estimate (β=0.533, 95% CI [0.271, 0.898], p = 0.027), despite a multimodal eulachon run in 2016 that contained three distinct peaks. The area under the curve eDNA rate explained 92.53% of the deviance in the mark-recapture population estimate (β=0.502, 95% CI [0.338, 0.697], p = 0.005) (Fig. 4). The peak eDNA concentration without flow correction explained 89.53% of the deviance in the mark-recapture population estimate (β=0.503, 95% CI [0.310, 0.742], p = 0.01). The area under the curve even without flow correction still explained 90.74% of the deviance in the mark-recapture population estimate (β=0.443, 95% CI [0.292, 0.620], p = 0.006) (Fig. 4). The quasipoisson regression models using either the flow-corrected eDNA rate peak (i.e. maximum of flow x DNA concentration) or the area under the curve as a single predictor produced highly representative predictions of mark-recapture population estimates (Fig. 5).

**Figure 3.**
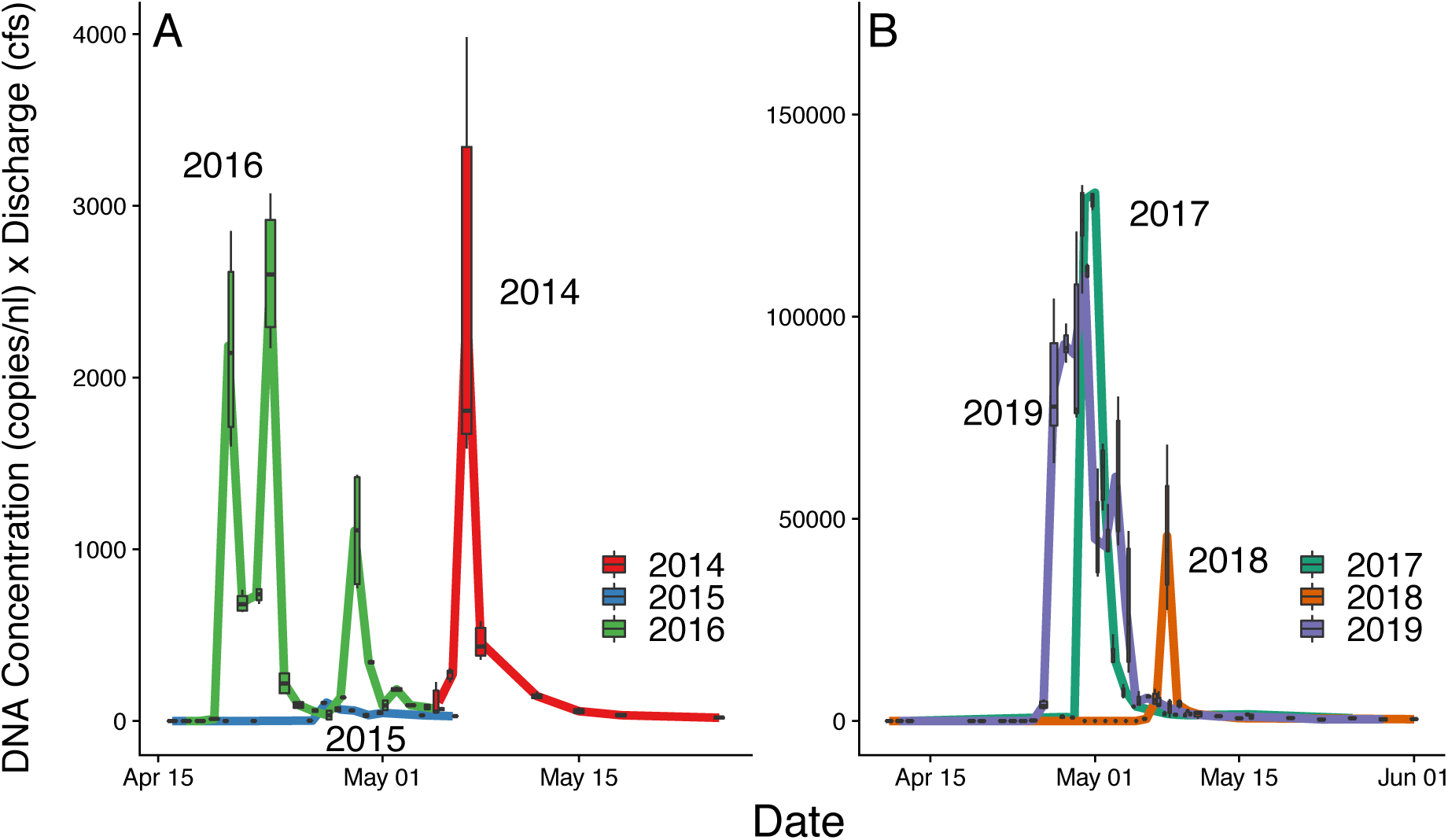
Daily results of the Chilkoot River flow-corrected eDNA rate (copies/nl * discharge in cubic-feet/sec.) in 2014-2016 (A), 2017-2019 (B). The boxplots illustrate the variability among the three daily water samples, each quantified in two ddPCR replicates. The lower and upper portions of the box correspond to the 25^th^ and 75^th^ quartile respectively around the median (line). Whiskers extend to the most extreme data points up to 1.5 times the interquartile range. Variability increased with the mean such that boxplots at low flow-corrected eDNA rates are very small and appear as points.

**Figure 4.**
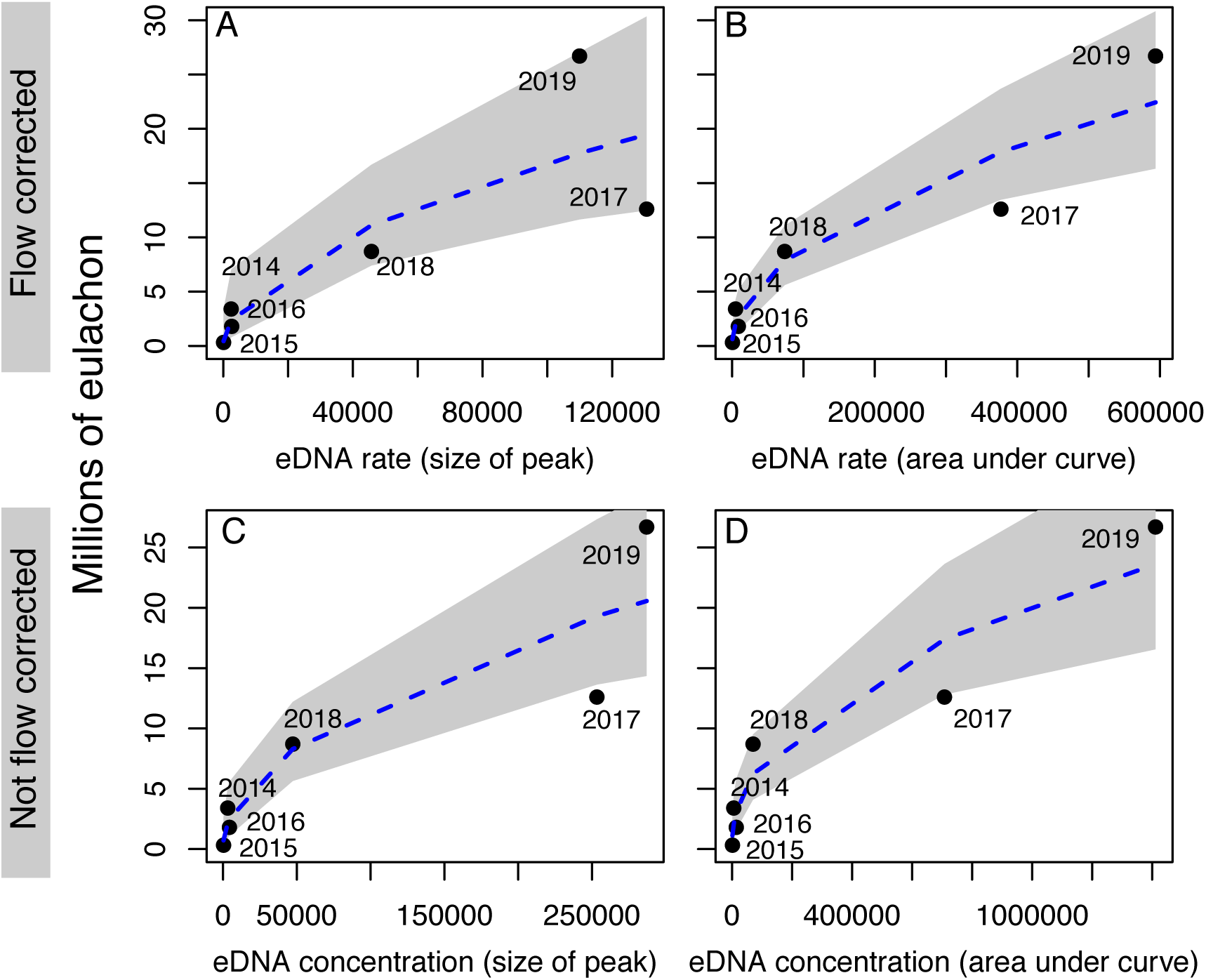
Results of quasipoisson regression models relating log-transformed mark-recapture population estimate to (A) the size of the peak flow-corrected eDNA rate (*p* = 0.027, 84.96% Deviance explained), (B) the area under the curve of the flow-corrected eDNA rate (*p* = 0.005, 92.53% Deviance explained), (C) the size of the peak uncorrected eDNA concentration (*p* = 0.01, 89.53% Deviance explained), and (D) the area under the curve of the uncorrected eDNA concentration (*p* = 0.006, 90.74% Deviance explained). Gray shading denotes 95% confidence interval.

**Figure 5.**
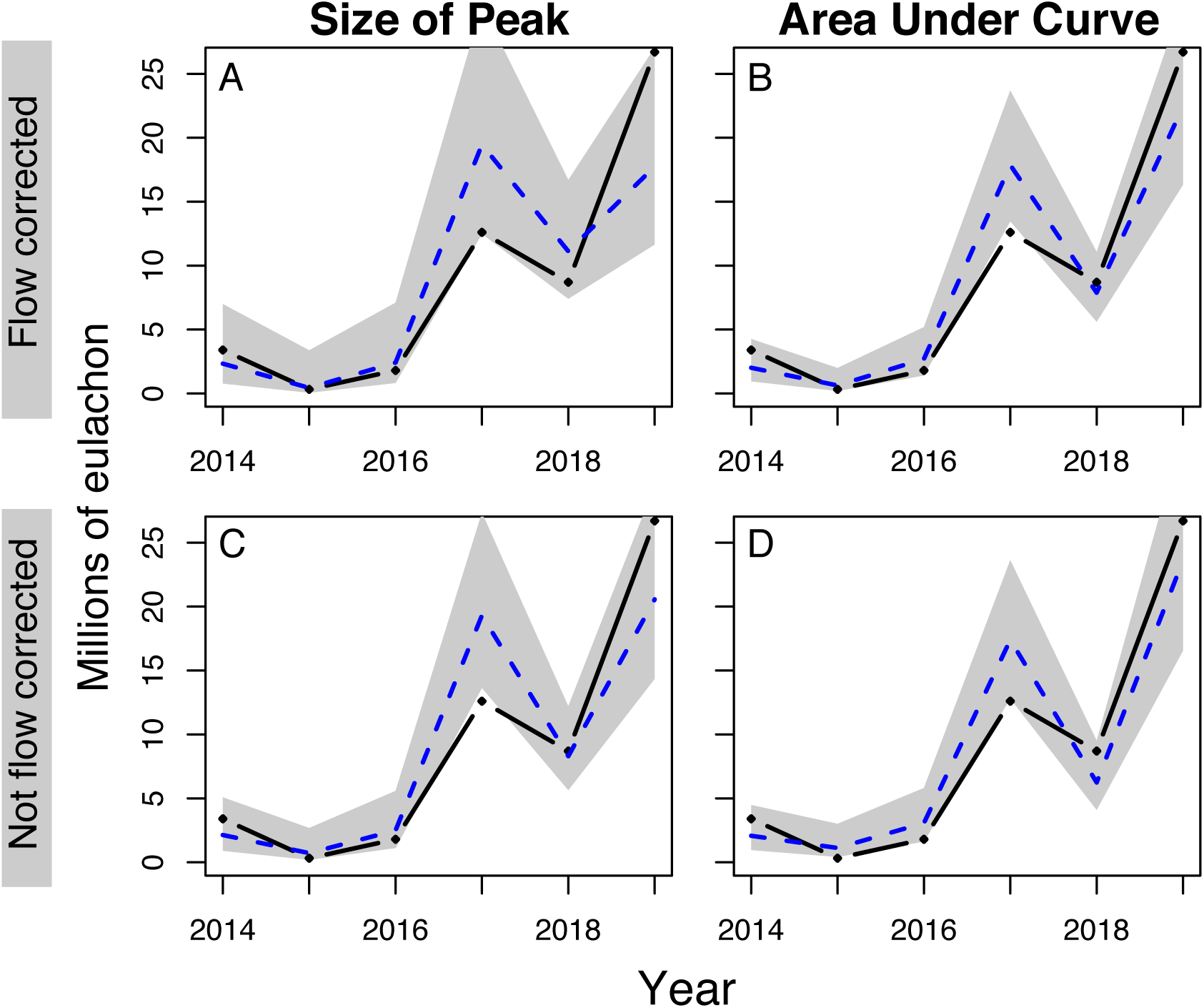
Mark recapture population estimate of eulachon runs (black dots) from 2014 to 2019 and the predicted number of eulachon based on the peak (A) or area under the curve (B) of the flow-corrected eDNA rate and the peak (C) and area under the curve (D) of the non-flow corrected eDNA concentration in the quasipoisson regression model (blue dashed lines). Gray shading denotes the 95% confidence interval.

## Discussion

The utility of eDNA for the detection of organisms has been widely documented (Ficetola, Miaud, Pompanon, & Taberlet, 2008; Rees et al., 2014; Wilcox et al., 2016). Recently, the next generation of eDNA science has evaluated the efficacy of quantifying the abundance of species using eDNA (Doi et al., 2015; Levi et al., 2019; Takahara et al., 2012; Tillotson et al., 2018). However, the expansion of eDNA beyond academic settings and into species management and monitoring is just beginning. eDNA methods may be particularly promising for the management of neglected species such as eulachon. This is true even if eDNA provides less accurate or precise results than do traditional methods, because lower quality data from more streams could result in more robust management decision-making than higher quality data from just a few streams (Dowling et al., 2008), particularly for a fish that exhibits low site fidelity. In addition, for many taxa, knowing population trends is just as important as precisely enumerating abundance. However, our results suggest that this tradeoff is largely inconsequential; both the flow-corrected and non-flow-corrected eDNA rate was predictive of the eulachon mark-recapture population estimates at a small fraction of the cost. Further, eDNA was predictive of mark-recapture population estimates even without flow-correction (Fig. 4D), which suggests the possibility that eDNA-based quantitation of eulachon could be implemented in systems where flow measurements cannot be obtained.

Unlike the mark-recapture method, which produced a single population estimate for the eulachon run, the eulachon eDNA rate captured within-run phenology as eulachon abundance varied in the Chilkoot River above the sampling location. eDNA was very effective at quantifying run timing and was particularly effective at demonstrating that the 2016 eulachon run was multimodal with three distinct pulses of eulachon that were separated by 4-5 days of inactivity (Fig. 3). The third pulse in 2016 was not represented in the mark-recapture estimate because field personnel had assumed that the run had terminated, but, due to the minimal labor required for eDNA sampling, we continued sampling and were able to capture the full 2016 run with eDNA.

Given the results presented here, eDNA has great potential as an indigenous-led, efficient, and effective means of monitoring this culturally and ecologically important species. Because eulachon in many regions, like northern Southeast Alaska, are not a commercially harvested species, they do not receive agency funding for management. This leaves a financial gap in monitoring that is often filled through indigenous-led efforts. A primary benefit of using eDNA is the vastly reduced cost of monitoring under-funded species, such as eulachon. The mark-recapture study on the Chilkoot River costs approximately $28,000 per year; largely because two 5-person crews are needed to properly implement the mark-recapture method. The use of eDNA at the current Oregon State University rate of $42/sample for 3 samples/day for ∼13 days is $3,000. Further cost savings would be accrued by reducing sampling prior to and after the run, when DNA concentrations are low, and instead focusing measurement during the ∼1 week peak of the run, because both the peak eDNA concentration and area under the curve eDNA metrics are invariant to sampling additional days with very low DNA concentrations. The economic viability of eDNA could be further increased with automated water samplers, currently in development, or the use of citizen scientists (Biggs et al., 2015; Wilken, 2018).

An additional benefit of eDNA methods is that mark-recapture estimation is not logistically feasible on all rivers. The Chilkoot poses a unique set of characteristics – single channel, road accessible, and with a relatively distinct upper limit to spawning activity. Many rivers in Southeast Alaska where eulachon spawn are glacially-fed, with wide, braided river mouths that are in remote, road-less areas. A mark-recapture method at these locations would be logistically challenging, in large part due to a large field-crew requirement. The appeal of eDNA is the ability to simply collect and filter a water sample to derive an index of abundance, which can be done by a single person in under one hour. The use of eDNA allows population data to be gathered on rivers that otherwise would not be possible, which is vital in monitoring a population that exhibits a regional genetic population structure where a decline in spawning biomass in any one river system might not necessarily represent a decline in the regional eulachon population (Flannery et al., 2013). The use of affordable and logistically feasible eDNA methods could facilitate regional studies of eulachon population size, run timing, and synchrony among rivers, which would allow for inference on regional population trends, environmental drivers of population dynamics, and environmental drivers of spawning river selection (Bryant, 2009).

Measurement of eDNA concentrations at a point in space and time represents a simple sampling process of mtDNA molecule counts per unit reaction volume, which can be modelled by a Poisson distribution assuming that eDNA is well mixed. The maximum likelihood estimate of the actual concentration of eDNA, λ, is equal to the sample mean of the *N* replicate eDNA concentrations, and the variance around this estimate is equal to the sample mean of the eDNA concentrations divided by *N.* Thus, the variance of the estimated ‘true’ eDNA concentration declines quickly with the number of replicates from a maximum equal to the mean. In contrast, mark-recapture analysis is a relatively complex statistical sampling process with a variance around the population estimate of ([(*M* + 1)(*C* + 1)(*M* − *R*)(*C* − *R*)]/[(*R* + 1)^2^(*R* + 2)]), which can be much larger than the mean population estimate (Table 1; SE^2^ can be thousands of times larger than N) if the number of recaptured individuals, *R*, is small relative to the number of marked individuals, *M*, or the number captured in the second session, *C* (Chapman, 1951). Thus, although eDNA concentrations are not in the useful units of individual animals, they can be estimated precisely with limited replication, but the same is not true for mark-recapture population estimates. Of course, while the observation error of eDNA concentrations in water samples may be low, the process error linking eDNA concentrations to fish abundance can be quite high due to the complexity of eDNA transport and degradation, variance in eDNA production among individuals and through time, the random spatial location of organisms relative to the sampling site, and more challenging stream morphologies among other complexities.

Although there is substantial process error linking eDNA concentrations to fish abundance, mark-recapture presents its own suite of problems. For example, the demographic-closure assumptions of mark-recapture estimators are difficult to meet with an anadromous fish that quickly enters and leaves the river (Pollock, 2018). The Chilkoot River mark-recapture study lasts for the duration of the run (typically 4-8 days), beginning on the first day that fish are observed in the river (typically late April) and ending once recapture sampling has exhausted all new fish into the system (i.e. when recaptures are identifying double-marked fish). During this time, new fish immigrate into the river while subsistence fishing activities actively remove fish, thus violating closure. However, mark-recapture population estimates can be robust to moderate violations of closure (Kendall, 1999). In this study, the closed-population assumption is thought to be reasonably met because (1) initial marking efforts remained relatively constant and continued until no new fish appear to be entering the system (i.e. approximately all individuals were potentially subject to marking), (2) there was an equal probability of capture of marked and unmarked fish by subsistence harvesters, and (3) we secured participation of subsistence harvesters to search their catch for marked fish.

We have demonstrated that eDNA provides reliable quantification of anadromous eulachon abundance. eDNA is thus a promising tool that can be mobilized by indigenous people to affordably monitor noncommercial species that are neglected by agencies but are culturally and/or economically important to native people. However, substantial uncertainty persists. In particular, it is unknown whether our model correlating eulachon run size with flow-corrected eDNA will be transferable to other rivers. This is unlikely to be the case when rivers have different morphologies, such as braided floodplains with pockets of eulachon spawning throughout. In such circumstances, a within-river index of abundance might be achieved by monitoring several braids where eulachon congregate, or perhaps the estuary where mixing of water might homogenize the sample. Once sampling specifications for rivers of various morphologies are determined, the final step toward large-scale implementation is regional cooperation among Alaska natives, First Nations in British Columbia, and tribal governments in Washington and Oregon to facilitate monitoring of eulachon spawning rivers throughout the Pacific Northwest.

## Acknowledgements

We thank Oregon State University, The National Geographic Society (#9493-14), US Fish & Wildlife Service Tribal Wildlife grant (AK-U-28-NA-1), North Pacific Landscape Conservation Cooperative (FW13AP01047), and Bureau of Indian Affairs (A18AP00229) for funding this work. Additionally, we thank the Chilkoot Indian Association Eulachon Tribal Working Group for incorporating the cultural significance of eulachon into this research and all the tribal members that worked as part of the mark-recapture crews. D.W. Yu was supported by the National Natural Science Foundation of China (41661144002, 31670536, 31400470, 31500305), the Key Research Program of Frontier Sciences, CAS (QYZDY-SSW-SMC024), the Bureau of International Cooperation (GJHZ1754), the Strategic Priority Research Program of the Chinese Academy of Sciences (XDA20050202, XDB31000000), the Ministry of Science and Technology of China (2012FY110800), the State Key Laboratory of Genetic Resources and Evolution (GREKF18-04) at the Kunming Institute of Zoology, and the University of East Anglia.

## Data Accessibility Statement

All data and code are made available in the supporting information.

## Author Contributions

TH and MP organized the eulachon mark-recapture program. TL initiated the eDNA monitoring program based on discussion with DY and SM. TL, MP, JA, collected eDNA samples that were processed in the lab by JA and MP. JA developed primers. MP and TL analyzed data and wrote the paper with input and edits from other authors.

## Notes

#### Summary of Updates

We have (1) made the framing more general, (2) provided more methodological detail on the eDNA assay, (3) included 2019 data, making it a 6-year time series, and (4) remedied an error. In big eulachon years, we diluted samples to avoid droplet saturation. This was properly accounted for in the flow-corrected model (ConcXflow variable), but not in the model that did not correct for flow (i.e. the models used Concentration instead of Conc_X_Dilution in supporting data). After correction, both flow-corrected and not flow-corrected models are good predictors of eulachon mark-recapture estimates.

## References

Barnes, M. A., & Turner, C. R. (2016). The ecology of environmental DNA and implications for conservation genetics. Conservation Genetics. https://doi.org/10.1007/s10592-015-0775-4

Betts, M. F. (1994). The subsistence hooligan fishery of the Chilkat and Chilkoot Rivers. Technical Paper Series, (213), 1–69.

Biggs, J., Ewald, N., Valentini, A., Gaboriaud, C., Dejean, T., Griffiths, R. A., … Dunn, F. (2015). Using eDNA to develop a national citizen science-based monitoring programme for the great crested newt (Triturus cristatus). Biological Conservation, 183, 19–28. https://doi.org/10.1016/J.BIOCON.2014.11.029

Bryant, M. D. (2009). Global climate change and potential effects on Pacific salmonids in freshwater ecosystems of southeast Alaska. Climatic Change, 95(1–2), 169–193. https://doi.org/10.1007/s10584-008-9530-x

Clarke, A. D., Lewis, A., Telmer, K. H., & Shrimpton, J. M. (2007). Life history and age at maturity of an anadromous smelt, the eulachon Thaleichthys pacificus (Richardson). Journal of Fish Biology, 71(5), 1479–1493. https://doi.org/10.1111/j.1095-8649.2007.01618.x

Doi, H., Uchii, K., Takahara, T., Matsuhashi, S., Yamanaka, H., & Minamoto, T. (2015). Use of Droplet Digital PCR for Estimation of Fish Abundance and Biomass in Environmental DNA Surveys. PLOS ONE, 10(3), e0122763. https://doi.org/10.1371/journal.pone.0122763

Dowling, N. A., Smith, D. C., Knuckey, I., Smith, A. D. M., Domaschenz, P., Patterson, H. M., & Whitelaw, W. (2008). Developing harvest strategies for low-value and data-poor fisheries: Case studies from three Australian fisheries. Fisheries Research, 94(3), 380–390. https://doi.org/10.1016/J.FISHRES.2008.09.033

Ficetola, G. F., Miaud, C., Pompanon, F., & Taberlet, P. (2008). Species detection using environmental DNA from water samples. Biology Letters, 4(4), 423–425. https://doi.org/10.1098/rsbl.2008.0118

Flannery, B. G., Spangler, R. E., Norcross, B. L., Lewis, C. J., & Wenburg, J. K. (2013). Microsatellite Analysis of Population Structure in Alaska Eulachon with Application to Mixed-Stock Analysis. Transactions of the American Fisheries Society, 142(4), 1036–1048. https://doi.org/10.1080/00028487.2013.790841

Hart, J. (1973). Pacific fishes of Canada. Bulletin of the Fisheries Research Board of Canada, 180, 148–150.

Hay, D., & Mccarter, P. B. (2000a). Status of Eulachon Thaleichtheys pacificus in Canada. Fisheries and Oceans Canada. Canadian Stock Assessment, 2000/145.

Hay, D., & Mccarter, P. B. (2000b). Status of eulachon Thaleichthys pacificus in Canada. Fisheries and Oceans Canada. Canadian Stock Assessment, 9R.

Jerde, C. L., Mahon, A. R., Chadderton, W. L., & Lodge, D. M. (2011). “Sight-unseen” detection of rare aquatic species using environmental DNA. Conservation Letters. https://doi.org/10.1111/j.1755-263X.2010.00158.x

Kendall, W. (1999). Robustness of Closed Capture – Recapture Methods To Violations of the Closure Assumption. Ecology, 80(8), 2517–2525.

Lacoursière-Roussel, A., Côté, G., Leclerc, V., & Bernatchez, L. (2016). Quantifying relative fish abundance with eDNA: a promising tool for fisheries management. Journal of Applied Ecology. https://doi.org/10.1111/1365-2664.12598

Laramie, M. B., Pilliod, D. S., & Goldberg, C. S. (2015). Characterizing the distribution of an endangered salmonid using environmental DNA analysis. Biological Conservation. https://doi.org/10.1016/j.biocon.2014.11.025

Levi, T., Allen, J. M., Bell, D., Joyce, J., Russell, J. R., Tallmon, D. A., … Yu, D. W. (2019). Environmental DNA for the enumeration and management of Pacific salmon. Molecular Ecology Resources. https://doi.org/10.1101/394445

Lincoln, F. C. (1930). Calculating waterfowl abundance on the basis of banding returns. US Department of Agriculture, 118, 1–4.

Lodge, D. M., Turner, C. R., Jerde, C. L., Barnes, M. A., Chadderton, L., Egan, S. P., … Pfrender Michael E. (2012). Conservation in a cup of water: estimating biodiversity and population abundance from environmental DNA. Molecular Ecology, 21(11), 2555–2558. https://doi.org/10.1111/j.1365-294X.2012.05600.x

Mächler, E., Deiner, K., Steinmann, P., & Altermatt, F. (2014). Utility of environmental DNA for monitoring rare and indicator macroinvertebrate species. Freshwater Science. https://doi.org/10.1086/678128

Moody, M. F., & Pitcher, T. J. (2010). Eulachon (Thaleichthys pacificus): past and present. The Fisheries Centre, University of British Columbia, Canada, 18(2), 197. https://doi.org/10.14288/1.0074735

NOAA. (2010). Endangered and Threatened Wildlife and Plants: Threatened Status for Southern Distinct Population Segment of Eulachon. In Federal Register (Vol. 75). https://doi.org/10.1021/j100299a032

Olds, A. (2016). Integrating local and traditional knowledge and historical sources to characterize run timing and abundance of eulachon in the Chilkat and Chilkoot Rivers. University of Alaska Fairbanks.

Petersen, C. G. J. (1896). The yearly immigration of young plaice into the Limfjord from the German Sea. Report of the Danish Biological Station, 6, 1–77.

Plough Id, L. V, Ogburn, M. B., Fitzgerald, C. L., Geranio, R., Marafino, G. A., & Richie, K. D. (2018). Environmental DNA analysis of river herring in Chesapeake Bay: A powerful tool for monitoring threatened keystone species. https://doi.org/10.1371/journal.pone.0205578

Pollock, K. H. (2018). Modeling Capture, Recapture and Removal Statistics for Estimation of Demographic Parameters From Fish and Wildlife Populations: Past, Present and Future. American Statistical Association, 86(413), 225–238.

Rees, H. C., Maddison, B. C., Middleditch, D. J., Patmore, J. R. M., & Gough, K. C. (2014). REVIEW: The detection of aquatic animal species using environmental DNA - a review of eDNA as a survey tool in ecology. Journal of Applied Ecology, 51(5), 1450–1459. https://doi.org/10.1111/1365-2664.12306

Sigler, M. F., Womble, J. N., & Vollenweider, J. J. (2004). Availability to Steller sea lions (Eumetopias jubatus) of a seasonal prey resource: a prespawning aggregation of eulachon (Thaleichthys pacificus). Canadian Journal of Fisheries and Aquatic Sciences, 61(8), 1475–1484. https://doi.org/10.1139/f04-086

Signorell, A. (2019). DescTools: Tools for description statistics.

Sowa, J. (ADFG). (2015). Hydrologic Investigations in Support of Reservations of Water for the Lost River, Alaska by.

Spangler, E. A. K. (2002). The Ecology of Eulachon (Thaleichthys pacificus) in Twentymile River, Alaska. Univeristy of Alaska Fairbanks.

Takahara, T., Minamoto, T., Yamanaka, H., Doi, H., & Kawabata, Z. (2012). Estimation of Fish Biomass Using Environmental DNA. PLoS ONE, 7(4), e35868. https://doi.org/10.1371/journal.pone.0035868

Team, Rs. (2015). RStudio: Integrated Development for R. Retrieved from http://www.rstudio.com

Tillotson, M. D., Kelly, R. P., Duda, J. J., Hoy, M., Kralj, J., & Quinn, T. P. (2018). Concentrations of environmental DNA (eDNA) reflect spawning salmon abundance at fine spatial and temporal scales. https://doi.org/10.1016/j.biocon.2018.01.030

Turnipseed, D. P., & Sauer, V. B. (2010). Discharge Measurements at Gaging Stations. Retrieved from http://pubs.usgs.gov/tm/tm3-a8/

Wilcox, T. M., McKelvey, K. S., Young, M. K., Sepulveda, A. J., Shepard, B. B., Jane, S. F., … Schwartz, M. K. (2016). Understanding environmental DNA detection probabilities: A case study using a stream-dwelling char Salvelinus fontinalis. Biological Conservation, 194, 209–216. https://doi.org/10.1016/j.biocon.2015.12.023

Wilken, U. (2018). Lakes, Labs and Learning: How an Environmental DNA Citizen Science Project… K-12 STEM Education, 4(4), 391–399. Retrieved from https://www.learntechlib.org/p/209557/

Yates, M., Fraser, D., & Derry, A. (2019). Meta-analysis supports further refinement of eDNA for monitoring aquatic species-specific abundance in nature. Environmental DNA, 1, 5–13. https://doi.org/10.1002/edn3.7

